# Interactions between parasites within the host: helminth infection reduces lice abundance in feral pigeons (*Columba livia*)

**DOI:** 10.1101/2025.02.25.640130

**Authors:** Aurélie Jeantet, Fabienne Audebert, Simon Agostini, Beatriz Decencière, Maxence Chéreau, Maïa Grasset, Pierre Federici, David Rozen-Rechels, Julien Gasparini

## Abstract

1. Within a host, parasites form a community and interact with each another. They can interact directly through competition for space or indirectly through competition for resources, or even via the host’s immune response. These interactions can influence infection patterns within the host. In particular, helminths are capable of facilitating the infection of other parasites through their immunosuppressive effects or hindering it by activating immune responses that reduce the ability of other species to infest the host.
2. In this study, we aimed to test the existence of parasite interactions in the wild pigeon host (*Columba livia*). More specifically, we experimentally tested whether helminth infestation influences the infestation dynamics of ecto- and endoparasites in pigeons.
3. Our predictions were tested on a sample of pigeons captured in Paris, half of them were exposed to an anthelmintic treatment. We then assessed the effects of this deworming treatment on the abundance of blood parasites (hemosporidian parasites, *Haemoproteus* spp., *Plasmodium* spp., and *Leucocytozoon* spp.), lice (*Columbicola columbae* and *Campanulote compar*), as well as on the intensity of antiparasitic behavioral responses (preening).
4. We detected a negative relationship between helminth presence and lice abundance, with higher lice abundance when helminths were reduced (with anthelmintic treatment) and lower lice abundance when helminths were more abundant (without anthelmintic treatment), indicating inhibitory effects of helminths on ectoparasite abundance. However, these results were not supported by differences in preening activity between the two experimental groups, suggesting that other mechanisms are involved. Our results showed no effect of anthelmintic treatment on the abundance of blood parasites. The mechanism by which helminths induced ectoparasite repulsion remains to be demonstrated.
5. Our study highlights the importance of considering interactions between different types of parasites in an eco-epidemiological approach when taking into account the factors affecting parasite prevalence and abundance.

## INTRODUCTION

Parasitism is defined as a long-term biological relationship between two organisms, the parasite and the host, where the parasite extracts energy from the host, using its host as a food source, habitat or for its own reproduction (Allan et al., 2020; Combes et al., 2018; Seppälä et al., 2008; Zelmer, 1998). Thus, this interaction can lead to a decrease in the host’s fitness, due to the direct and indirect costs imposed by the parasite (Koop et al., 2011). These costs may include energy losses induced by parasites and defenses activated by the host to fight against parasites. For example, these defenses can take the form of costly immune or behavioral antiparasitic responses (Ezenwa, 2004; Gray et al., 2012; Zuk & Stoehr, 2002). The activation of such defenses may indeed impact in the escape capacities, the fertility or even the growth of hosts, which can then lead to the decline of the population dynamics of the hosts (Allan et al., 2020; Marcogliese, 2004). From an ultimate point of view, the parasites significantly impact the fitness of the host and constrain the evolution of the host. According to Price (1980), every living organism is affected by parasitism, either as a host, or as a parasite, or both and many scientists also consider that more than 50% of animal species are parasites, or have at least one parasitic phase in their life cycle (Lucius et al., 2017; Windsor, 1998).

There exists a wide variety of parasites, which are found in various locations on their host’s body. Parasites found on the surface of the host’s body are called ectoparasites (Hopla et al., 1994). An example of ectoparasites is lice, such as *Haematomyzus elephantis* which infests elephants, or flies, such as *Carnus hemapterus* which parasitizes nesting birds as adults, as well as a wide variety of crustaceans, mosquitoes, ticks, etc. which can be specific to a host species or infest a wide range of hosts species (Hopla et al., 1994). Parasites living inside the body of their host, with access to the external environment, are called mesoparasites. Among them, we find intestinal parasites, which live in the digestive tract of their host, such as *Toxocara cati* in felines, or *Ancylostoma duodenale* in hoofed mammals (Lim et al., 2008). Finally, parasites living inside the body of their host, without access to the external environment, are called endoparasites. They feed on fluids, as do blood cell parasites. For example, *Haemoproteus* spp. in birds, transmitted by flat flies, but also *Trichomonas gallinae* or *Plasmodium* sp. which is transmitted by mosquitoes (Alkharigy et al., 2018; Atkinson et al., 2008).

Thus, within their host, these parasites form a community and interact with each other (Biard et al., 2015; Graham, 2008). They can interact directly through competition for space (interference competition), or indirectly through competition for resources (bottom-up interaction, exploitation competition) or the host immune response (top-down interaction) (Biard et al., 2015; Graham, 2008; Nunn et al., 2014). These interactions between parasites can influence infection patterns at the host level. In particular, studies have shown that helminths are able to modulate host infestation by other parasite species (ecto-, endo- and mesoparasites). First, they can facilitate the infection of other parasites through their immunosuppressive effects (“immune response cost” hypothesis, Graham, 2008; Nunn et al., 2014). For example, Graham (2008) showed by a meta-analysis that helminths infecting mice would induce an increase in the abundance of endoparasites when they do not share the same resources, via the suppression of a cytokine (interferon gamma (IFN)-?), involved in the defense against endoparasites.

Second, other studies have shown that helminths can also compete with other parasites when they share the same resources or by activating immune responses that will reduce the ability of other species to infest the host (parasite competition hypothesis, Larsen et al., 2002). This is what Larsen et al. (2002) demonstrated, showing that infection by helminths (*Anisakis* sp.) would decrease the abundance of ectoparasites (*Gyrodactylus derjavin*) in trout (*Salmo trutta*) compared to uninfected fish. This phenomenon would be due to the activation of the immune system of trout infected by intestinal worms, which would then modify the host’s skin response and create a negative effect on ectoparasites.

Altogether, these studies have shown the ability of mesoparasites to modulate the infestation capacities of other parasites, which would be partly mediated by positive or negative effects on the antiparasitic defenses of the hosts. Furthermore, these effects of mesoparasites on the infestation dynamics by other parasites depend on the species of parasites and hosts considered.

In this study, we aim to test the existence of interactions between parasites in feral pigeon host (*Columba livia*). We experimentally tested whether mesoparasite infestation influences ecto- and endoparasite infestation dynamics. If the “immune response cost” hypothesis prevails (Nunn et al., 2014), we expected that helminth infection would facilitate infection by Haemosporidian parasites. Alternatively, if the “parasite competition hypothesis” prevails, we predicted that helminth infection reduces the intensity of ectoparasites (Larsen et al., 2002). As we expected that this interaction might be mediated by the impact of mesoparasites on host antiparasitic behaviors, we also tested whether the presence of mesoparasites influenced preening activity, an effective strategy against plumage ectoparasites.

## MATERIAL AND METHODS

For this study, we used a subsample of pigeons (n = 66) non-exposed to lead exposure of a previous study testing another hypothesis (Jeantet et al., 2024). The sampling and the general protocol of anthelmintic treatment are reported in Jeantet et al. (2024). Hereafter, we briefly summarize the bird sampling and the anthelmintic treatment protocol.

### Bird sampling

A sample of 66 feral pigeons was captured in January 2022 in different Parisian districts. To identify the pigeons individually, we put a colored and numbered ring on their left paw and a colored ring on their right paw. After their capture, birds were placed in 6 outdoor aviaries (dimensions: 3 x 2.2 x 2.2 m) at the Foljuif station in Saint-Pierre-lès-Nemours, France (Center for Research in Experimental and Predictive Ecology, CEREEP UMS 3194, 48° 17’N, 2° 41’ E). They were distributed in a way to equilibrate their melanic coloration (Kruskall-Wallis, χ^2^_5_ =0.61, P-value = 0.99), their sex (GLM, χ^2^_5_ =3.11, P-value = 0.68), their mass (ANOVA, F_5, 60_ = 0.20, P-value = 0.96) and their site of capture (Fisher’s exact test, P-value = 0.98), in order to limit experimental biases and confounding effects linked to their past life history.

### Experimental protocol

All pigeons were placed in their aviaries at least two weeks before the start of the experimental treatment, in order to acclimatize them to their new environment. At the start of the experiment, the aviaries were divided into two experimental groups: (1) without anthelmintic treatment (presence of helminths, n=33), (2) with anthelmintic treatment (absence of helminths, n=33). For the anthelmintic group, we administered 600 μL of a solution containing levamisole (Capizol 15 mg/mL) on the first day of the experimental treatment and every month after. This anthelmintic treatment was administered to the birds directly in the beak using a syringe, and in concentrations considered effective in the literature for the elimination of nematodes (Chege et al., 2017). In order to ensure that all birds were subjected to the same handling stress, birds without anthelmintic treatment (presence of mesoparasites) received an equivalent volume of water, using the same method and at a similar frequency. At the end of the experiment, we confirmed the effectiveness of the anthelmintic treatment through coproscopies, following the methodology described in Jeantet et al. (2024). To achieve this, we employed the MacMaster flotation technique. We first collected the feces from each pigeon then weighed and floated in a 15 mL Falcon tube containing a saturated NaCl solution. The tubes were shaken, and after a 15-minute settling period, we collected the supernatant and applied it to a MacMaster slide.

This method allowed us to count worm eggs under a microscope, enabling us to compare the nematode abundance between treated and untreated pigeons. As a result, we identified the presence of several taxa: *Capillaria* sp., *Syngamus trachea, Ascaridia* sp. and *Trichostrongylus tenuis*. Furthermore, the results revealed that the total abundance of nematodes was significantly higher in pigeons without anthelmintic treatment, compared with pigeons with anthelmintic treatment (with anthelmintic: 7.87 helminth eggs / g of feces ± 3.57, without anthelmintic: 39.9 helminth eggs / g of feces ± 8.35; χ^2^_1_ =8.18, P<0.01), indicating that our anthelmintic treatment was effective. Finally, during all the experiment, pigeons were fed ad libitum with peas, maize and wheat and aviaries were enriched with baths and bamboo sticks as perches. At the end of the experiment, all birds were released near their capture sites.

### Endoparasite abundance

In order to measure the abundance of Hemosporidian parasites (*Heamoproteus* spp., *Plasmodium* spp., *Leucocytozoon* spp.), blood smears were realized the first week of the experiment and at weeks 5, 9, 13, 17, 21, 26. Blood smears were fixed in methanol and stained with May-Grünwald Giemsa coloration (Sordolab kit, ref COLRASA), to reveal blood parasites. All smears were then photographed using an optical microscope equipped with a camera and the number of infected blood cells over 5000 blood cells were counted using ImageJ software.

### Ectoparasite abundance

Quantification of the abundance of ectoparasites *Columbicola columbae* and *Campanulotes compar* was carried out by visual examination according to the method described by (Koop & Clayton, 2013), at weeks 10 and 24. In order to exert the same pressure on all pigeons, a standardized 5-minute protocol was established. This protocol began with the examination of the right wing then the left wing for a duration of 1 minute 30 per wing, followed by the observation of the neck, head and body for 1 minute 30 also, before finishing with the observation of the tail for 30 seconds. In order to limit bias, all birds were examined by the same observer. In addition, each part of the birds’ body was observed only once in order to avoid counting the same lice several times. Thus, the variable used to quantify ectoparasite abundance was the number of *Columbicola columbae* and *Campanulotes compar* counted in the plumage of pigeons.

### Preening activity

In order to quantitatively measure grooming activity, behavioral scans were performed at weeks 7, 15, 16, 18, 20, 22 and 24. Each pigeon was observed for 30 minutes during weeks 6 and 8 (15 minutes in the morning and 15 minutes in the afternoon) and 15 minutes in the other weeks, bringing the total observation time of each pigeon to 165min. To limit bias, the order in which the aviaries were observed as well as the observers varied between each scan. Thus, the variable used to quantify preening activity was the mean number of preening behaviors observed during each behavioral scan.

### Statistics

Statistical analyses were performed using R (version 4.1.2). To test the effect of anthelminthic treatment on ecto- and endoparasite abundances and preening, we ran three models in which we systematically added anthelmintic treatment, time (expressed in week as a categorical variable) and their interaction. We also added in all our models the pigeon ID nested in the aviary as a random effect and sex as a cofactor fixed effect. To test the effect of anthelmintic treatment on ecto- and endoparasite abundances, we used two generalized linear mixed models (GLMM) for negative binomial distribution (glm.nb function from the MASS package), in order to control overdispersion. For endoparasites, we also included the initial blood parasite abundance as covariates. Finally, to test the effect of helminth removal on preening, we also ran a GLMM for negative binomial distribution.

## RESULTS

### Ectoparasite abundance

We detected a significant effect of the interaction between anthelmintic treatment and time (χ^2^_1_ = 7.6, P= 0.006, Fig. 1). At week 10, lice abundance did not differ between pigeons with anthelmintic treatment and control pigeons (χ^2^_1_ = 0.09, P = 0.77). However, at week 24, lice abundance was significantly higher in pigeons with anthelmintic treatment than in control pigeons (χ^2^_1_ = 6.04, P = 0.01). Moreover, we detected a significant effect of sex on lice abundance (χ^2^_1_ = 6.74, P = 0.009), with a higher abundance in male than in female (18.06 lice ± 2.91 SE in males and 11.21 lice ± 1.26 SE in females).

**Figure 1.**
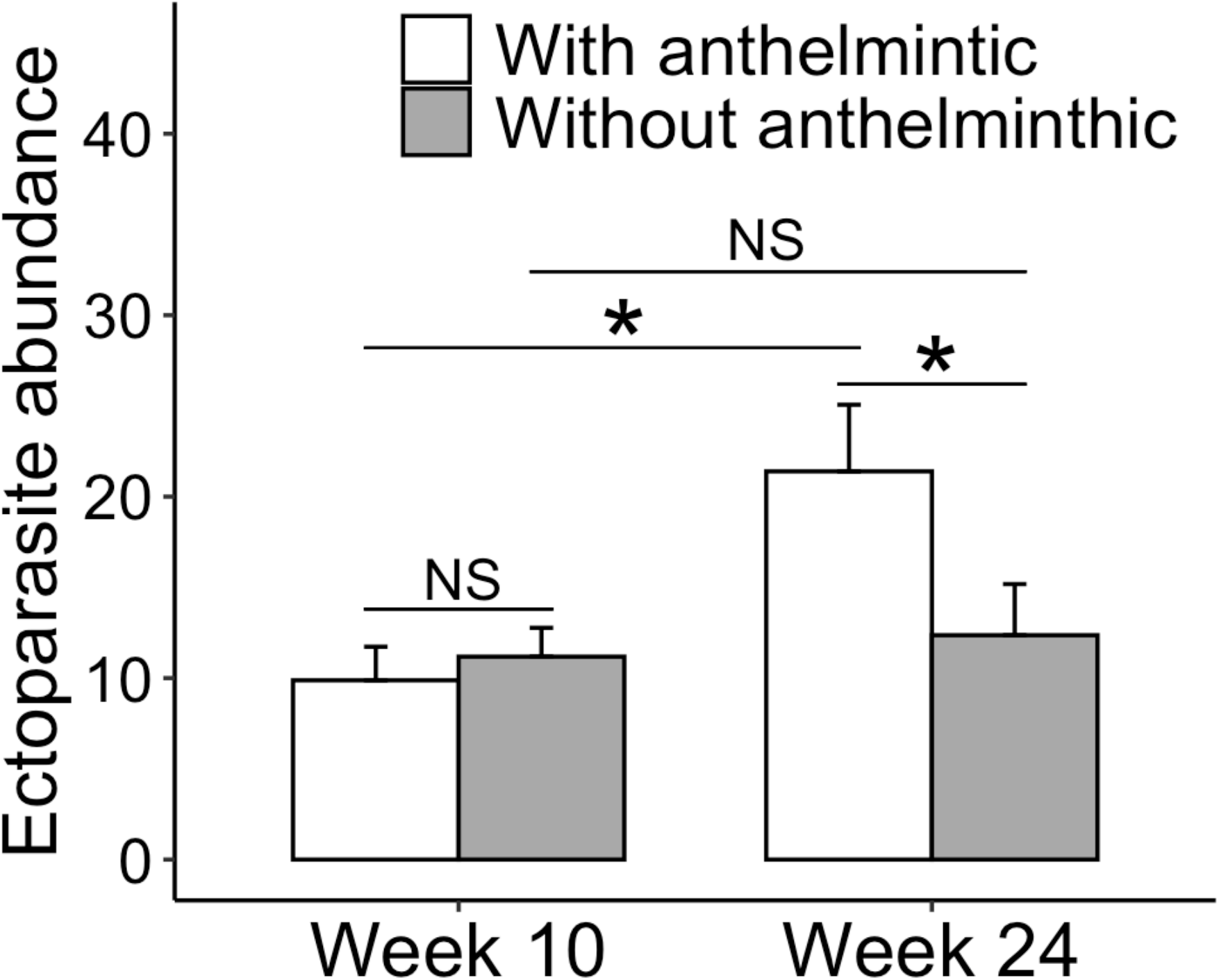
Ectoparasite abundance (*Columbicola columbae* and *Campanulotes compar*, mean + SE) across weeks 10 and 24, in birds with anthelmintic treatment (white bars) or without anthelmintic treatment (grey bars). Significant difference between groups is indicated by an asterisk (NS: p > 0.05, *: p< 0.05). SE, standard error.

### Endoparasite abundance

We did not detect any effect of anthelmintic treatment alone (χ^2^_1_ = 0.005, P=0.94), or in interaction with time (χ^2^_6_ = 3.29, P=0.77) on blood parasites. However, we detected a significant effect of time (χ^2^_1_ = 33.22, P<0.001).

### Preening activity

We detected a significant interaction between anthelmintic treatment and time on preening activity (χ^2^_8_ = 17.94, P = 0.02, Fig. 2). Indeed, at week 22, we detected a significant difference between groups, with a higher preening activity in pigeons with anthelmintic treatment than in control pigeons (χ^2^_1_ = 6.67, p = 0.01).

**Figure 2.**
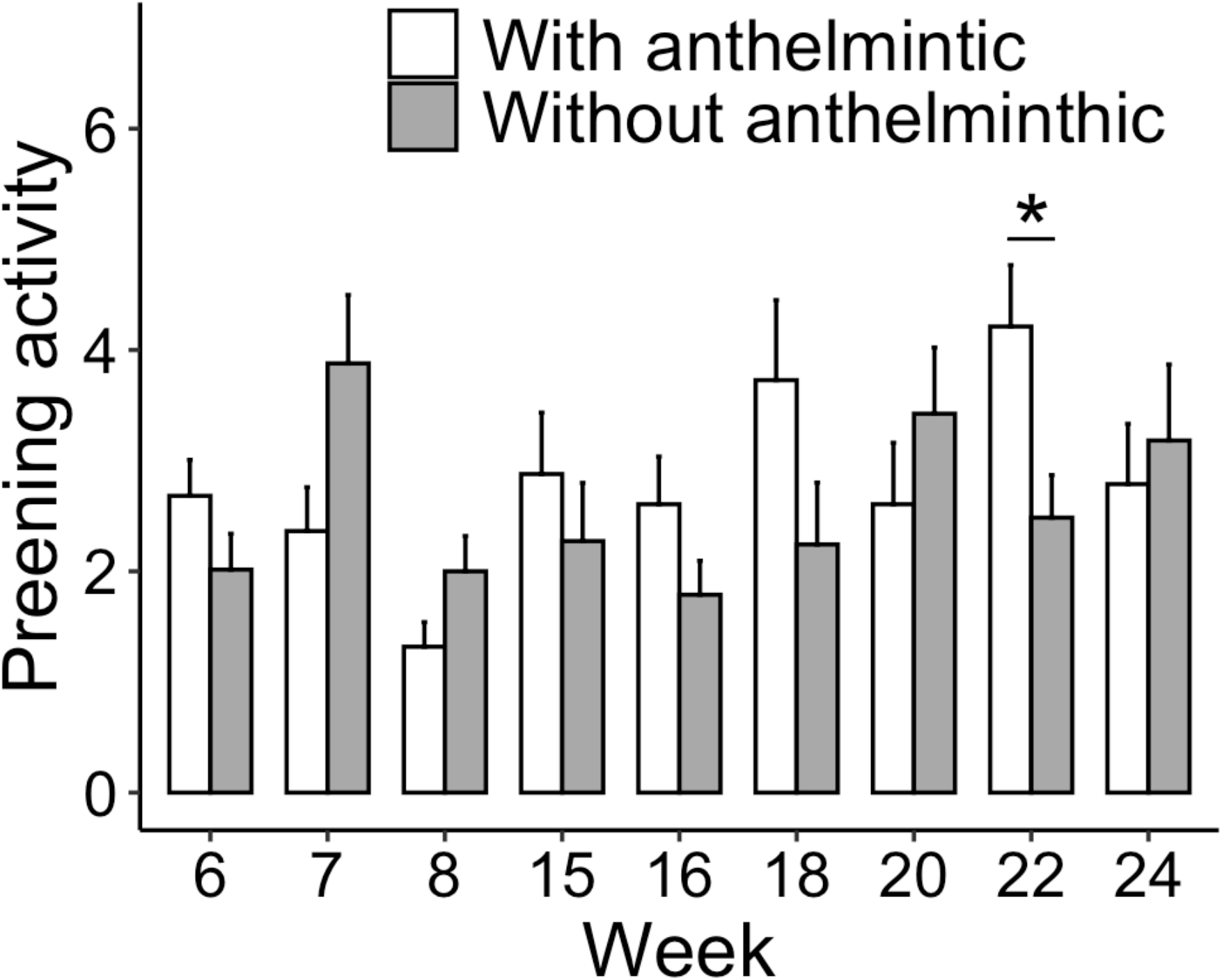
Preening activity (number of times each bird preened their feathers during each behavioral scan session, mean + SE) across 9 weeks, among birds with anthelmintic treatment (white bars) or without anthelmintic treatment (grey bars). Significant difference between groups is indicated by an asterisk (NS: p > 0.05, *: p< 0.05). SE, standard error.

## DISCUSSION

Our results are consistent with the hypothesis that it exists interactions between parasites within an individual in feral pigeon. Indeed, we found an effect of anthelmintic treatment on ectoparasite abundance, with a higher abundance when helminths are reduced (with anthelmintic treatment) than with higher abundance of helminths (control). This significant interaction reveals a negative effect of helminths on ectoparasites. In contrast, we did not observe any effects of anthelmintic treatment on blood parasite abundance (Fig. 2). Furthermore, contrary to our prediction, our results do not suggest that the presence of helminths increases preening activity, which would have explained the potential negative effects of helminths on lice abundance (which is not the case in this study).

First, our results do not support the immune cost hypothesis (Holmstad et al., 2008), as the anthelmintic treatment did not significantly impair endo- and ectoparasite abundances. According to (Graham, 2008), mesoparasites would induce a decrease in the immunocompetence of hosts, which could facilitate infection by endoparasites. In agreement with our result, our previous study did not show any effects of anthelmintic treatment on the immune response (Jeantet et al., 2024) which, in turn, did not affect endoparasites. Similarly, in this study, we did not find a strong effect of anthelmintic treatment on the preening activity of pigeons. Indeed, we observed at week 22 more preening in pigeons without helminths. The cost associated with this behavioral anti-ectoparasites is not strongly reduced by the presence mesoparasites.

These results are not in agreement with those of (Saumier et al., 1988) who observed a reduction in grooming activity in American Kestrels (*Falco sparverius*) experimentally infected with a roundworm (*Trichinella pseudospirallis*). We believe that performing scan samplings rather than focal samplings over short periods of time may explain the low variability observed between individuals. It would have been interesting to carry out focal samplings instead, in order to detect finer differences between groups without helminths and with helminths. Finally, it would be interesting to carry out other experiments to investigate by which pathway this regulation of ectoparasites is carried out. If the hypothesis that the positive effect of helminths on endoparasites would be mediated by immune pathways is true (physiological of behavioral), it is possible that the lack of significant effects is linked to the ad libitum feeding provided to the pigeons throughout the experiment. Indeed, indirect cost of parasite effects may be amplified under low-resource conditions, possibly due to the increased immune costs associated with lowresource conditions (Bize et al., 2010). The negative impact of mesoparasites on ecto- and endoparasites, on preening activity in the present study and on the PHA immune response (Jeantet et al., 2024) may have been hidden by the high-resource condition of our experiment (food ad libitum).

Second, our results are rather consistent with the competition hypothesis as we detected a negative effect of helminths infection on the abundance of ectoparasites (Figure 1). This confirms the results of Larsen et al. (2002), who observed a reduction in the infection levels of ectoparasites (*Gyrodactylus derjavini*) in trout fry (Salmo trutta) experimentally infected with mesoparasites (*Anisakis* sp.). Larsen et al. (2002) hypothesized that the reduction in ectoparasite infection levels may result from skin reactions triggered by the encystment of helminth larvae. This process would activate the fish immune system in response to the mesoparasites, subsequently exerting negative effects on the ectoparasites. It would be interesting to investigate whether similar mechanisms occur in pigeons.

In conclusion, our experimental results support our general hypothesis, suggesting that interactions between parasites do occur in feral pigeon, but only between mesoparasites and ectoparasites. Interestingly, it seems that mesoparasites infection impaired infection by ectoparasites, suggesting an existence of competition between these two types of parasites. However, the mechanism by which helminth induced repulsion of ectoparasites remain to be identified. More generally, our study underlines the fact of considering interactions among the different types of parasites in eco-epidemiological approach when the factors affecting prevalence and abundances of parasites.

## ACKNOWLEDGEMENTS

This work was supported by the French National program EC2CO (Ecosphère Continentale et Côtière). AJ’ PhD grant was funded by by the ‘Biodiversity, Evolution, Ecology, Society’ initiative of the Sorbonne University Alliance. This study was carried out in strict accordance with the recommendations of the European Convention for the Protection of Vertebrate Animals used for Experimental and Other Scientific Purposes (revised Appendix A). All experiments and captures were approved by Charles Darwin Animal Experimentation Ethics Committee and French authorities (the “Ministère de l’éducation nationale, de l’enseignement supérieur et de la recherche”, permit N°#17554 2018111610466351 ; and Ville de Paris).

## AUTHOR CONTRIBUTIONS

Aurélie Jeantet, Julien Gasparini and Fabienne Audebert conceived the ideas and designed the methodology; Aurélie Jeantet analyzed the data and led the writing of the manuscript together with Julien Gasparini, David Rozen-Rechels and Fabienne Audebert. All the authors collected the data, contributed critically to the drafts and gave final approval for publication.

## DATA AVAILABILITY STATEMENT

The data will be deposited in Zenodo or other equivalent archives when accepted.

